# ClearX9-Stem™: an efficient, affordable & sustainable stem cell culture medium for biotech and cell-cultivated meat industries

**DOI:** 10.1101/2023.05.24.542098

**Authors:** Sumit Gautam, Neeraj Verma, Siddharth Manvati, Pawan K. Dhar

## Abstract

Stem cells are extraordinary cells with a unique ability of self-renewal and differentiation into various cell types such as muscle, nerve, bone, and blood cells. Historically, they have found significant applications in the biotech and pharma sectors. To grow and maintain stem cells artificially, researchers use basal media formulations supplemented with nutrients and growth factors, with Fetal bovine serum (FBS) as the key component of the culture medium. However, to maintain this supply every year, millions of pregnant cows are slaughtered for preparing FBS. The process of harvesting FBS also raises concerns about contamination with pathogens, animal proteins that may interfere with cellular behavior and ethical considerations regarding animal welfare. To overcome these limitations, here we report ClearX9-Stem™ - an affordable, sustainable, effective, and ethical replacement for an FBS-enriched stem cell culture medium. A specialized ClearX9-Stem™ cell culture medium formulation was designed to grow chicken embryonic fibroblast (SL-29) in the absence of FBS. Based on the results obtained, ClearX9-Stem™ is undergoing further refinement to meet the growing academic and industrial demand for serum-free culture media formulations. In the future, there is a need to customize and optimize ClearX9-Stem™ for the scalable growth of cells in bioreactors.

**HIGHLIGHTS:**

- ClearX9-Stem™ provides good nutritional support for the growth of chicken embryonic fibroblast cells.
- ClearX9-Stem™ cell growth performance is comparable to the serum-enriched culture medium
- ClearX9-Stem™ maintains a healthy morphological profile of cells during division
- ClearX9-Stem™ generates a stress-free environment within cells
- ClearX9-Stem™ does not require animal slaughter and reduces the environmental footprint
- ClearX9-Stem™ has applications in the biotechnology, pharma, and cell-cultivated meat industries

## 1. INTRODUCTION

Stem cells are unique cells with a remarkable ability of self-renewal and differentiation into various specialized cell types in the body. They serve as the building blocks of development and tissue regeneration. There are two primary types of stem cells: (i) Embryonic Stem Cells (ESCs) derived from embryos that are typically a few days old, usually obtained from in-vitro fertilization clinics or donated embryos. ESCs are pluripotent i.e., they have the capacity to differentiate into any cell type in the body. Adult Stem Cells (also known as Somatic or Tissue-specific Stem Cells) are present in specific tissues and organs throughout the body and serve as a repair and maintenance system for the body, replenishing damaged or aging cells. In addition, induced pluripotent stem cells (iPSCs), have been generated in the laboratory by reprogramming adult cells (such as skin cells) back into a pluripotent state.

Stem cells are of great interest to scientists and researchers due to their regenerative potential and ability to give rise to specialized cells. They hold promise for various applications in medicine, including tissue engineering, regenerative therapies, drug discovery, and disease modeling. Stem cell research continues to advance our understanding of developmental biology and offers potential solutions for treating diseases, injuries, and degenerative conditions.

Mesenchymal stem cells (MSCs) are self-renewing cells that can give rise to various mesodermally-derived tissues (Caplan and Bruder, 2001; Pittenger et al., 1999). Due to their multipotency, self-renewal capacity, and immunomodulatory properties, MSCs have attracted significant attention in regenerative medicine and other research fields (Nakamura et al., 2008).

One of the most critical unmet needs of the lab-grown meat, biotech, and pharma industries is the high cost of stem cell culture media for large-scale culture. For example, two tissue-engineering companies, Organogenesis (Canton, MA) and Advanced Tissue Sciences (La Jolla, CA), filed for bankruptcy due to enormous operating costs (Bouchie, 2002, 2003). There is a pressing need for effective alternatives that can sustain the supply, bring down the costs and meet the concerns of pathogen infection.

The cultivated meat industry uses FBS-enriched media to grow skeletal muscle stem cells (i.e., myosatellite cells), fibroblasts, mesenchymal stem cells, induced pluripotent, embryonic stem cells, and adipose-derived cells. However, FBS comes with several limitations like batch-to-batch variability, the risk of transmitting pathogens to humans, and the slaughter of pregnant cows leading to immense suffering and environmental costs (Gautam et al.; Tuomisto and Teixeira De Mattos, 2011).

To address pressing environmental, ethical, and safety concerns, ClearX9-Stem™ was designed as a natural growth media to provide nutrients for stress-free chicken embryonic fibroblast (SL-29) cell growth in-vitro.

## 2. MATERIAL AND METHODS

### 2.1 Cell line and reagents

Chicken Embryonic Fibroblast; SL-29 (CRL-1590™) cell line were obtained from American Type Culture Collection (ATCC) (Manassas, VA) and grown in Dulbecco’s Modified Eagle Medium (DMEM) (Invitrogen) and supplemented with 10% heat-inactivated fetal bovine serum (Gibco; Cat. No. 16170078) enriched cell culture medium or ATCC-formulated Dulbecco’s Modified Eagle’s Medium (DMEM), Catalog No. 30-2002. To make the complete growth medium, add the following components to the base medium: tryptose phosphate broth (TPB) (Hi-Media; SKU: M953) to a final concentration of 5% and fetal bovine serum to a final concentration of 5% and 1% penicillin-streptomycin (Invitrogen) or Clear X9-Stem™ at 5% CO2, 37°C under a humidified chamber.

ClearX9-Stem™ was designed to offer nutrients and growth factors under natural and non-slaughter conditions. The MTT [3-(4,5-Dimethylthiazol-2-Yl)-2,5-Diphenyl-tetrazolium Bromide] dye (M2128) was obtained from Sigma Aldrich. Triple distilled water (Millipore) was used for preparing solutions. All reagents used in the experiments were of the highest purity grade.

### 2.2 MTT Assay

The impact of standard Fetal Bovine Serum (FBS) supplemented with 5% tryptose phosphate broth (TPB) and FBS-free ClearX9-Stem™ medium was assayed on the proliferation rate of SL-29 cells using MTT assay. Briefly, all cells (5× 10^4^/ml) were plated (∼10,000 / well) in triplicate in 96 well plates after the minimum three passages of the SL-29 cell line. The cells were incubated in the presence of 10% FBS-enriched traditional (DMEM) or 5% FBS with 5% TPB supplemented in ATCC-formulated DMEM, (Cat. No. 30-2002) cell culture medium or ClearX9-Stem™ in a final volume of 200 µl for 72 hours at 37°C in a 5% CO2 humidified chamber. Cells grown with 10% FBS-enriched traditional (DMEM) or 5% FBS with 5% TPB supplemented in ATCC-formulated DMEM, (Cat. No. 30-2002) cell culture medium served as a control in this experiment. At the end of the cell culture, 20 μl of MTT solution (5 mg/ml in 1X PBS) was added to each well and incubated for additional 5 hours. The MTT-containing medium was discarded. The resultant formazan crystals were dissolved in 200µl of DMSO. The absorbance value (A) was measured at 570 nm by a microplate ELISA reader.

### 2.3 Cell morphology study

SL-29 Cells (7× 10^5^/ml) were plated in 2 ml, 10% FBS-enriched traditional (DMEM) or 5% FBS with 5% TPB supplemented in ATCC-formulated DMEM, (Cat. No. 30-2002) cell culture medium served as a control in this experiment or ClearX9-Stem™ in 35 mm^2^ culture dishes for 72 hours at 5% CO2, 37°C under a humidified chamber after the minimum three passages of SL-29 cell line. Bright-field microscopy was used for studying morphology in response to the presence of containing traditional/conditional cell culture medium or ClearX9-Stem™. All images were captured at 20X magnification using a bright field microscope (Zeiss Primovert inverted microscope).

### 2.4 Statistical analysis

Statistical analyses were carried out using Graph Pad Prism software. Experimental data were expressed as means ± S.D. and the significance of differences was analyzed by a one-way ANOVA test. The Dunnett test for used for testing experimental groups against the control group concerning cell viability.

## 3. RESULTS

Cell viability of SL-29 (Chicken Embryonic Fibroblast) cells was examined using an MTT assay.

Results suggested that ClearX9-Stem™ significantly (p<0.001) increased the chicken embryonic fibroblast (SL-29) cell’s growth rate as compared to the cells grown in 10% Fetal Bovine Serum (FBS) or 5% Fetal Bovine Serum with 5% Tryptose Phosphate Broth (TPB) supplemented cell culture medium (Fig. 1 and 3). Cellular morphology, using vacuole concentration, has been traditionally used as a standard parameter for defining the health of a chicken embryonic fibroblast (SL-29) cell.

**Fig. 1.**
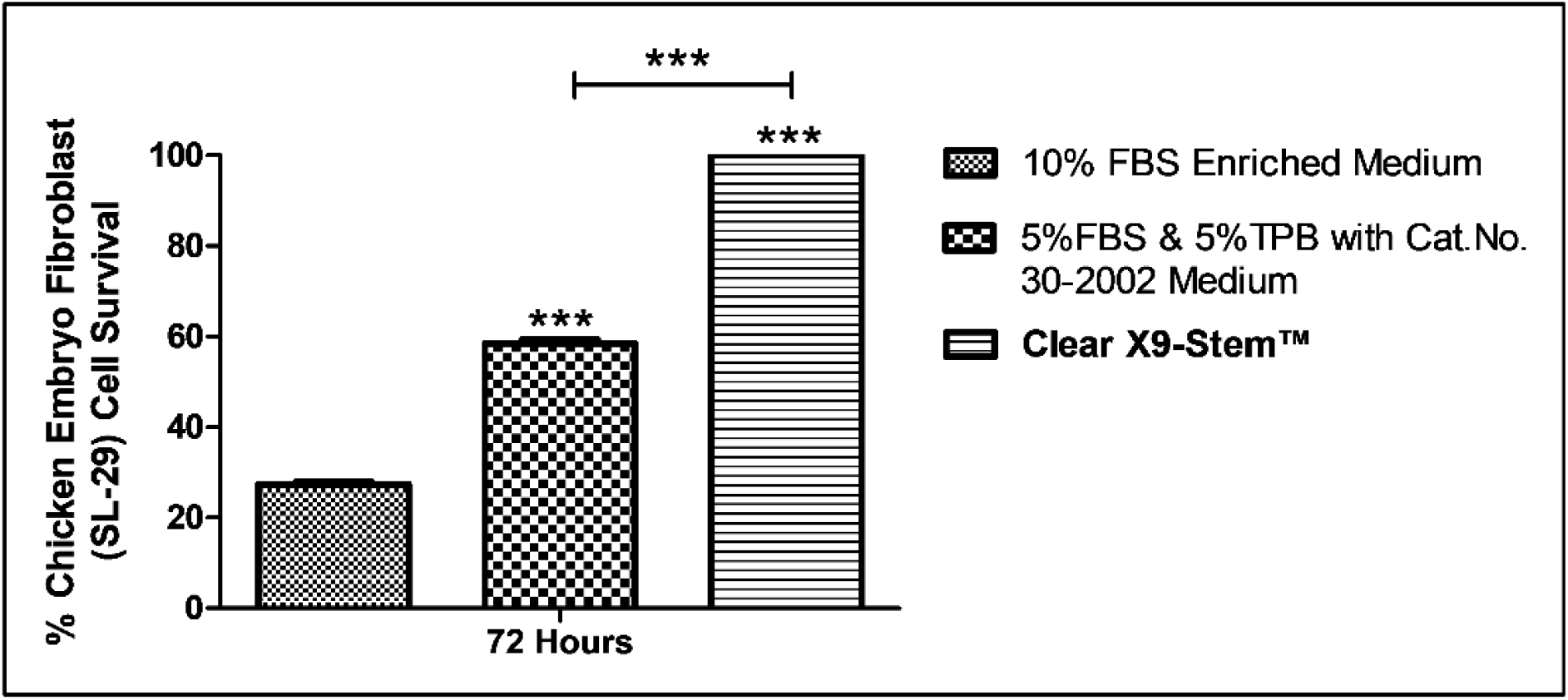
**ClearX9-Stem™ enhanced cells proliferation rate of SL-29 (Chicken Embryonic Fibroblast) cells at 72 hours. Cell Viability Assay was done by MTT on SL-29 cells. Briefly, cells (5 x104 cells/ml) were plated in 96 well plates and grown with 10% Fetal Bovine Serum (FBS) enriched containing standard (DMEM) or conditional (ATCC-Formulated DMEM (Cat. No. 30-2002) + 5% Tryptose Phosphate Broth + 5% Fetal Bovine Serum) complete medium or ClearX9-Stem™ for 72 hours. After the end of the stipulated time interval, MTT dye was added and incubated for 5 hours in a 37°C and 5% CO_2_ humidified chamber. The formed crystals were dissolved in DMSO and OD was taken at 570nM. The data shown are the mean from three parallel experiments, and each experiment was done in triplicate. ***p<0.001.**

Microscopic studies suggest a vacuole-free cytoplasmic presentation in cells growing in ClearX9-Stem™ as compared to the cells grown in 10% Fetal Bovine Serum (FBS) or 5% Fetal Bovine Serum with 5%

Tryptose Phosphate Broth (TPB) supplemented cell culture medium. A less clear cytoplasmic presentation was found in cells growing in FBS enriched medium indicating the presence of unhealthy cells as compared to the ClearX9-Stem™ cultured chicken embryonic fibroblast (SL-29) cells (Fig. 2 and 3).

**Fig. 2.**
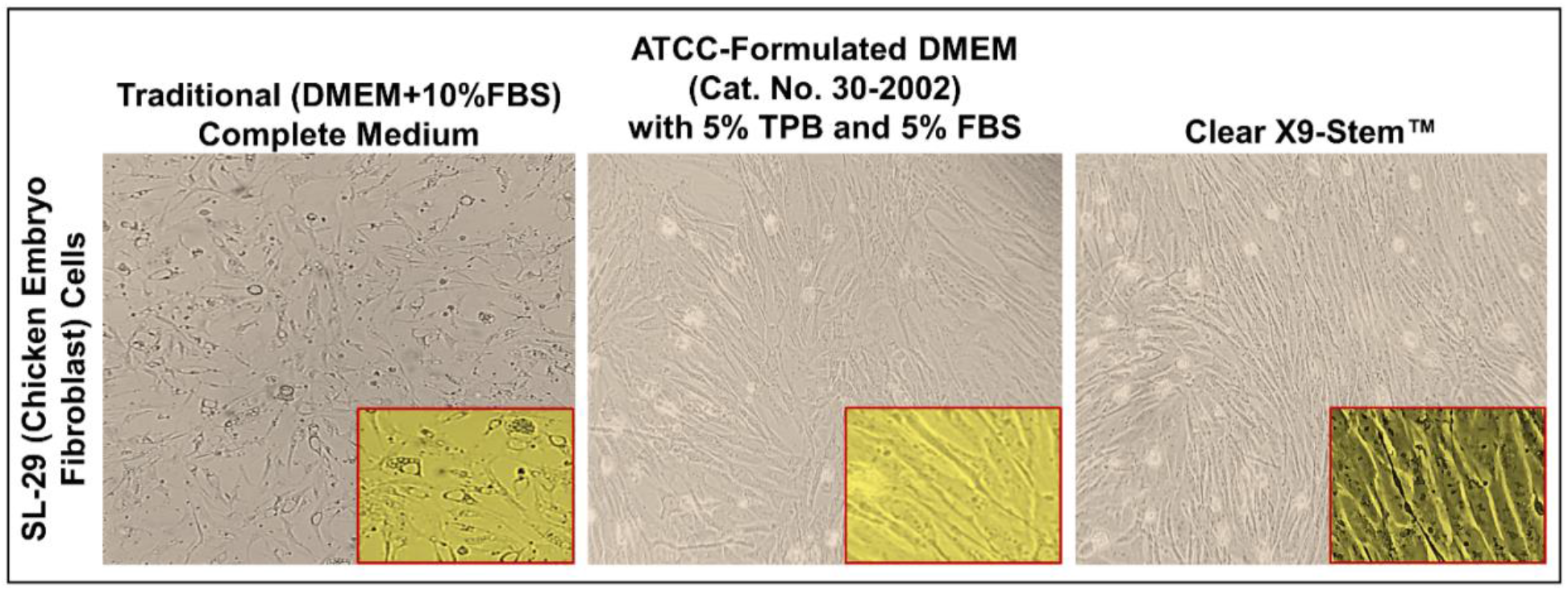
**ClearX9-Stem™ increased the SL-29 (Chicken Embryonic Fibroblast) cells growth rate at 72 hours. 7 × 105 cells/ml cells were plated in a 35mm2 culture dish and grown in the presence of 10% Fetal Bovine Serum (FBS) enriched containing standard (DMEM) or conditional (ATCC-Formulated DMEM (Cat. No. 30-2002) + 5% Tryptose Phosphate Broth + 5% Fetal Bovine Serum) complete medium or ClearX9-Stem™ and cell morphology was visualized after 72 hours using a bright field microscope (Images were captured by Zeiss Primovert inverted microscope at 20X magnification**

**Fig 3.**
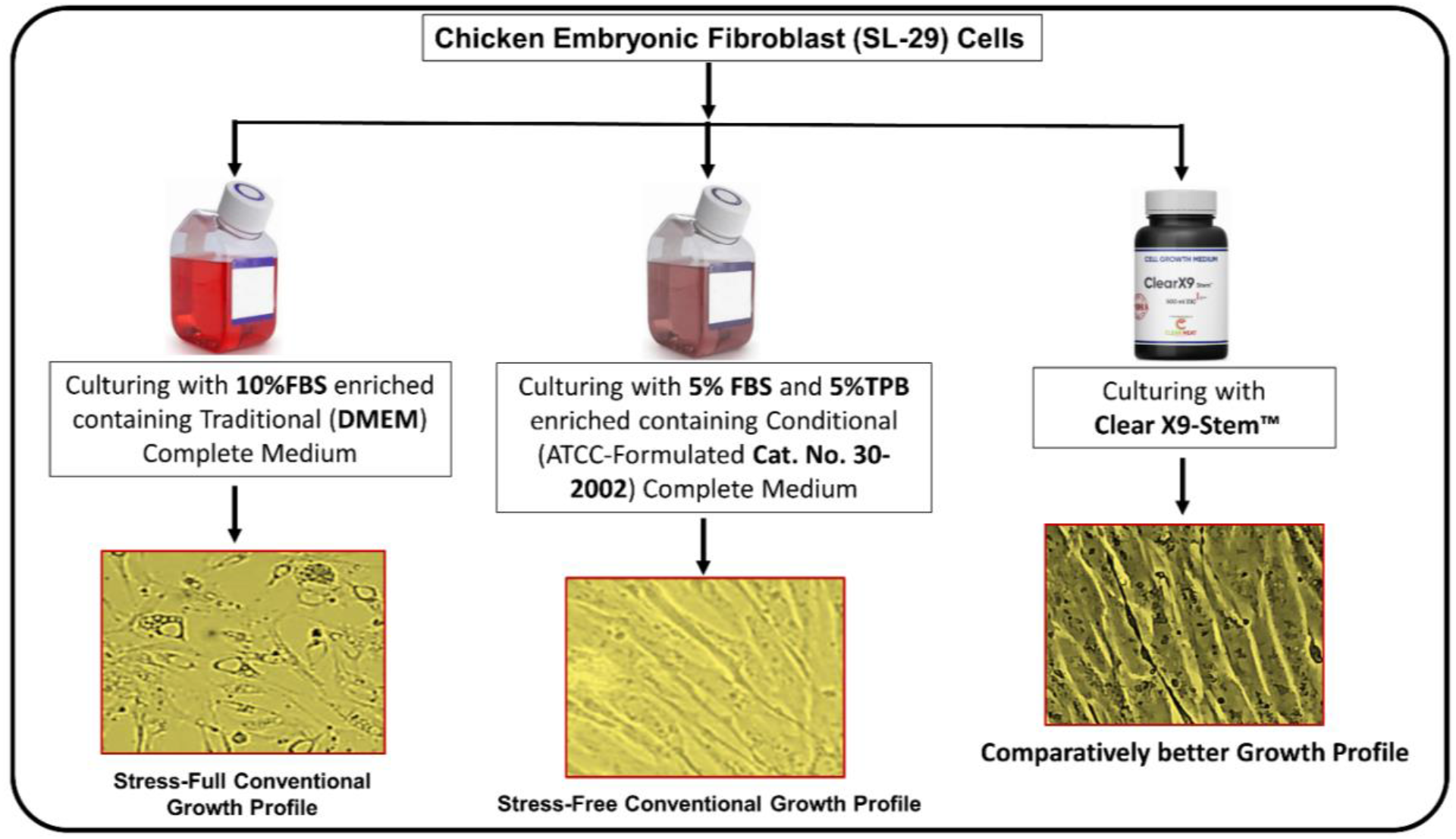
SL-29 (Chicken Embryonic Fibroblast) cells grown in ClearX9-Stem™ & Fetal Bovine Serum and Tryptose Phosphate Broth enriched cell culture medium. The conventional cell growth is clearly distinguished from the ClearX9-Stem™-supported cell growth

Results suggest a maximal cell growth rate in ClearX9-Stem™ compared to the cells grown in 10% Fetal Bovine Serum (FBS) or 5% Fetal Bovine Serum with 5% Tryptose Phosphate Broth (TPB) supplemented cell culture medium (Fig. 1 and 3). ClearX9-Stem™ showed a significantly (p<0.001) increased growth rate of SL-29 cells with features of good health and stress-free condition compared to 10% FBS-enriched traditional (DMEM) or 5% FBS with 5% TPB supplemented in ATCC-formulated DMEM, (Cat. No. 30-2002) cell culture medium cells (Fig. 2 and 3).

## 4. DISCUSSION

Cell culture media refer to the nutrient-rich formulations that support the growth, survival, and proliferation of cells in a laboratory setting. Culture media are designed to provide cells with the necessary nutrients, salts, vitamins, growth factors, and other essential components required for their growth and maintenance outside of their natural environment.

Cell culture media typically consist of a basal medium providing a balanced combination of inorganic salts, sugars, amino acids, and vitamins necessary for cell growth, at an optimal pH and osmotic condition to create an environment that mimics the natural state within the body.

The fast growth of the cell culture media market is mainly driven by the surging requirements in pharma, biotech and increasing investment in research (Culture et al., 2023). In 2022, the global cell culture media market size was valued at USD 4,075 million and was predicted to grow at a compound annual growth rate (CAGR) of 12.23% from 2023 to 2030.

The major component of a stem cell culture medium is a fetal bovine serum (FBS), which provides a range of nutrients, growth factors, and other biological components for supporting the growth, proliferation, and differentiation of cells. However, FBS comes with several limitations e.g., exorbitant cost, batch-to-batch variability, risk of pathogen transmission, and slaughter of pregnant cows leading to immense suffering to the animal.

Reports indicate that two million bovine fetuses are used worldwide on average to produce approximately 800,000 L of FBS. The non-stringent regulation of the FBS market results in the dilution of FBS quality, sustains cruelty to animals, generating huge methane emissions from the cattle industry (Sinke et al., 2023; Treich, 2021; Tuomisto and Teixeira de Mattos, 2010; Tuomisto and Teixeira De Mattos, 2011). The disadvantages can also arise from ill-defined descriptions of components, and the presence of adverse factors such as endotoxin, mycoplasma, viral contaminants, or prion proteins (Gstraunthaler et al., 2013). Due to this reason, there is a pressing need for innovative solutions.

These and many more concerns have accelerated the search for a credible alternative to FBS-enriched stem cell culture medium and built a new stem cell culture medium for the sustained growth of mesenchymal stem cells, induced pluripotent, embryonic stem cells, fibroblasts, starter cells, skeletal muscle stem cells (i.e., myosatellite cells) and adipose-derived cells.

Researchers have been exploring possibilities in the form of serum-free or serum-reduced media supplemented with growth factors, cytokines, or other additives, and plant-based or synthetic serum supplements to provide consistent and reproducible cell culture conditions while minimizing animal slaughter. However, efforts seem to be largely localized and lab-based with little impact on the global FBS market.

In this context, the development of ClearX9-Stem™ assumes significance, as ClearX9-Stem™ meets performance indicators of cell growth parameters in comparison to the cells grown in the FBS-enriched media. Cells cultured in ClearX9-Stem™ show clarity of cytoplasm indicating a stress-free environment compared to traditional/conditional complete medium condition (Fig. 3).

ClearX9-Stem™ has been designed to provide a ready-to-use, natural component-based product, with an adequate supply of amino acids, vitamins, lipids, plant pigments, and natural growth promoters for the growth of animal stem cells in-vitro.

In this report, we provide the first experimental evidence of using ClearX9-Stem™ to grow chicken embryonic fibroblast (SL-29) cells.

Results indicate that ClearX9-Stem™ mediated proliferation of SL-29 cells retains a healthy growth and a healthy morphology, compared to the cells grown in 10% Fetal Bovine Serum (FBS) or 5% Fetal Bovine Serum with 5% Tryptose Phosphate Broth (TPB) supplemented traditional/conditional cell culture medium cells (Fig. 3). Data strongly suggest ClearX9-Stem™ as a complete and sustainable stem cell culture medium for biotechnology and lab-grown meat industries.

Cellular morphology, using vacuole concentration, has been traditionally used as a standard parameter for defining the health of a cell. Microscopic studies suggest a vacuole-free cytoplasmic presentation in cells growing in ClearX9-Stem™ as compared to the Fetal Bovine Serum (FBS) enriched containing standard (DMEM) cell culture medium cells (Fig. 3). A less clear cytoplasmic presentation was found in cells growing in FBS and Tryptose Phosphate Broth enriched containing conditional (ATCC-Formulated DMEM (Cat.No. 30-2002) complete medium indicating the presence of unhealthy/dead cells (Crowley et al., 2016; Thong et al., 2016).

In chicken embryonic fibroblast (SL-29) cells, vacuoles were found to be present in 10%FBS enriched traditional (DMEM) complete medium indicating stressful conditions within cells in comparison to the stress-free cytoplasmic environment observed in ClearX9-Stem™ grown cells (Fig. 3).

This study suggests the potential of ClearX9-Stem™ as a credible FBS-enriched stem cell culture medium alternative in the biotech and cell-cultivated meat sectors. More work is required to study the application of ClearX9-Stem™ in areas like vaccine manufacturing, regenerative medicine, biomanufacturing, and toxicology.

Furthermore, there is a need to perform in-depth pathway analysis to answer cell growth and cell health questions at the molecular level. In the future, it would be interesting to know how genes are up/down regulated in FBS-enriched stem cell culture medium vs. ClearX9-Stem™ combinations.

## ACKNOWLEDGEMENT

Clear Meat Pvt. Ltd. expresses its sincere gratitude to key investors Gastrope Accelerator Group (supported by Miseltoe), BRINC Group, Artesian HB Group, and Gene Kim for funding this work.

## AUTHOR’S CONTRIBUTIONS

The concept and formulation of the ClearX9-Stem™ medium were conceived by PKD and supervised by SM for optimization. All the experiments were jointly performed by SG and NV. The manuscript was written by SG & PKD. All authors approved the data presented in this manuscript.

[**SG**: Sumit Gautam, **NV**: Neeraj Verma, **SM**: Siddharth Manvati, **PKD**: Pawan K. Dhar]

## Notes

### Competing Interest Statement

Sumit Gautam, Neeraj Verma are full time employees of Clear Meat Pvt Ltd. Siddharth Manvati works as CEO and Pawan K Dhar as Chief Mentor in the company. Both Siddharth Manvati and Pawan K Dhar own stocks in the company.

## REFERENCES

1. Bouchie, A. (2002). Tissue engineering firms go under. Nat. Biotechnol. 20.

2. Bouchie, A. (2003). Industry ponders reimbursement crisis. Nat. Biotechnol. 21.

3. Caplan, A.I., and Bruder, S.P. (2001). Mesenchymal stem cells: Building blocks for molecular medicine in the 21st century. Trends Mol. Med. 7.

4. Crowley, L.C., Marfell, B.J., Christensen, M.E., and Waterhouse, N.J. (2016). Measuring cell death by trypan blue uptake and light microscopy. Cold Spring Harb. Protoc. 2016.

5. Culture, C., Market, M., Analysis, T., By, R., Application, B., End-user, B., and Region, B. (2023). Report Overview. 1–14.

6. Gautam, S., Verma, N., Manvati, S., and Dhar, P.K. ClearX9 TM : an efficient alternative to fetal bovine serum for growing animal cells in vitro.

7. Gstraunthaler, G., Lindl, T., and Van Der Valk, J. (2013). A plea to reduce or replace fetal bovine serum in cell culture media. Cytotechnology 65, 791–793.

8. Nakamura, S., Yamada, Y., Baba, S., Kato, H., Kogami, H., Takao, M., Matsumoto, N., and Ueda, M. (2008). Culture medium study of human mesenchymal stem cells for practical use of tissue engineering and regenerative medicine. Biomed. Mater. Eng. 18.

9. Pittenger, M.F., Pittenger, M.F., Mackay, A.M., Mackay, A.M., Beck, S.C., Beck, S.C., Jaiswal, R.K., Jaiswal, R.K., Douglas, R., Douglas, R., et al. (1999). Multilineage potential of adult human mesenchymal stem cells. Science (80-.). 284.

10. Sinke, P., Swartz, E., Sanctorum, H., van der Giesen, C., and Odegard, I. (2023). Ex-ante life cycle assessment of commercial-scale cultivated meat production in 2030. Int. J. Life Cycle Assess. 28.

11. Thong, K.T., Ismail, A.B., Basri, H., Tee, K.S., and Soon, C.F. (2016). The effects of culture substrates and media on the behavior of microtissues. ARPN J. Eng. Appl. Sci. 11.

12. Treich, N. (2021). Cultured Meat: Promises and Challenges. Environ. Resour. Econ. 79.

13. Tuomisto, H.L., and Teixeira de Mattos, J.M. (2010). Life cycle assessment of cultured meat production. 7th Int. Conf. Life Cycle Assess. Agri-Food Sect.

14. Tuomisto, H.L., and Teixeira De Mattos, M.J. (2011). Environmental impacts of cultured meat production. Environ. Sci. Technol. 45.

